# Mapping eukaryotic chromatin accessibility and histone modifications with DNA deaminase

**DOI:** 10.1101/2024.12.24.630236

**Authors:** Wenyang Dong, Jiankun Zhang, Leyi Dai, Jinxin Phaedo Chen, Honggui Wu, Runsheng He, Yuxuan Pang, Zhi Wang, Fanchong Jian, Jun Ren, Yijun Liu, Yu Tian, Shuang Liu, Xuechen Zhao, Xiaoliang Sunney Xie

**Affiliations:** Changping Laboratory; Beijing, 102206, China; Beijing Advanced Innovation Center for Genomics (ICG) & Biomedical Pioneering Innovation Center (BIOPIC), Peking University; Beijing 100871, China; School of Life Sciences, Peking University; Beijing, 100871, China; Academy for Advanced Interdisciplinary Studies, Peking University; Beijing, 100871, China; College of Chemistry and Molecular Engineering, Peking University; Beijing, 100871, China

## Abstract

A variety of methods such as ATAC-seq and CUT&Tag have been developed for probing chromatin states. However, enzymatic cleavage of DNA in these methods hinders their multi-omics integration with other DNA profiling techniques. Here, by extending our use of dsDNA deaminase (FOOtprinting with DeamInasE, FOODIE), we introduce two new methods for mapping Chromatin Open region by DeamInasE (ChrODIE) and histone modification by ANtibody-associated DeamInasE (ANDIE), respectively. Unlike cleavage-based methods, both ChrODIE and ANDIE leave genomic DNA after deamination unfragmented and amplifiable. This offers the potential for simultaneous or downstream integration with other genomic profiling methods, such as mapping 3D genome interactions.

## Introduction

In eukaryotic cells, DNA is packaged into chromatin within the nucleus. The fundamental unit of chromatin organization is the nucleosome, and the organization of nucleosomes on DNA have a critical function in transcriptional regulation. Traditionally, a region of chromatin is said to be “open” or “accessible” if it is amenable to cleavage by a nuclease, typically an exogenous nuclease introduced for the purpose of mapping open regions. Areas of open chromatin correspond to non-nucleosomal areas of the genome that are typically bound by sequence-specific factors and their co-factors, many of which function to activate or repress gene transcription (1, 2). Several enzymatic cleavage-based methods, such as deoxyribonuclease hypersensitive site sequencing (DNase-seq) (3), micrococcal nuclease sequencing (MNase-seq) (4), and Assay for Transposase-Accessible Chromatin with high-throughput sequencing (ATAC-seq) (5), have been developed to map open chromatin or nucleosome positioning.

Covalent modification of histones can either promote or inhibit the recruitment of transcription factors and chromatin-associated enzymes, thereby modulating nucleosome stability, transcription, and other DNA-templated processes (6). Cleavage Under Targets and Tagmentation (CUT&Tag) and Cleavage Under Targets and Release Using Nuclease (CUT&Run) (7, 8) are methods that tether enzymes (either the Tn5 transposase used in ATAC-seq or MNase) to antibodies through fused protein A/G. This arrangement enables target-specific fragmentation or cleavage of genomic DNA, resulting in high-sensitivity high-resolution maps of histone modifications and chromatin-associated proteins.

However, all these techniques involve enzymatic cleavage of DNA, which prohibit many other simultaneous or downstream DNA profiling approaches. To address this, methods based on using DNA-modifying enzymes to probe chromatin accessibility have been developed. These include NOMe-seq, which uses cytosine methyltransferase M.CviPI and requires a bisulfite-conversion treatment to detect exogenous methylation by sequencing (4). More recently, DNA N6-adenine methyltransferase has enabling transcription factor footprinting (Fiber-seq) at high deaminase concentrations and open chromatin mapping (AdMTase-seq) at lower concentrations (9). Similar to CUT&Tag, the use of antibody-tethered N6-adenine methyltransferase in DiMeLo-seq allows for mapping histone modifications or chromatin-associated proteins based on targeted 6mA methylation. (10). However, 6mA modifications in Fiber-seq and DiMeLo-seq are not preserved during PCR amplification and therefore require third-generation sequencing (e.g. PacBio SMRT or Oxford Nanopore) for detection, posing a technical limitation.

Deaminases are members of a superfamily of enzymes that are widely distributed across eukaryotic and prokaryotic organisms, catalyzing the deamination of DNA (single- and double-stranded), RNA, nucleosides, and other nucleotide derivatives (11, 12). Deaminases are involved in processes such as immunity and metabolism (13, 14). Single-stranded DNA (ssDNA) deaminases, including enzymes that act on adenosine, cytidine, and guanine, have been utilized in various biotechnological applications, such as DNA-cleavage-free genome-editing techniques (e.g., base editors) (15, 16), or mapping epigenetic modifications in DNA and RNA (17, 18). One such example is the Cabernet technique based on the APOBEC deaminase, which enables single-cell bisulfite-free 5mC and 5hmC sequencing (19). Double-stranded cytosine deaminases have been recently exploited for use in mitochondrial DNA editing (12), or one-step enzymatic 5mC detection on dsDNA (14).

Rather than develop tools for gene editing, we aimed to repurpose dsDNA deaminase for studying chromatin and transcription. We previously developed FOODIE (FOOtprinting with DeamInasE), a novel technique enabling genome-wide identification of genomic sites bound by human transcription factors with near single-base resolution on a single-molecule and single-cell basis (20). In this study, we present a dsDNA deaminase-based epigenetic profiling strategy that directly maps Chromatin Open region by DeamInasE (ChrODIE) and histone modification by ANtibody-associated DeamInasE (ANDIE) on DNA molecules. Chromatin profiling with deaminase does not involve DNA cleavage and can be detected after PCR amplification, which holds the promise for integration with other genomic methods in a single assay.

## Results

### ChrODIE detects open chromatin regions

In the ChrODIE workflow (Fig. 1A), permeabilized nuclei are treated with DddA (Fig. 1B), a dsDNA deaminase that catalyzes cytosine (C) to uracil (U) conversion without DNA fragmentation. Following library preparation and sequencing, converted sites can be precisely identified by comparison to reference sequence. The cytosine deamination of genomic DNA by DddA acts only on cytosine preceded by thymine (TC) (Fig. 1C) (12).

**Figure 1:**
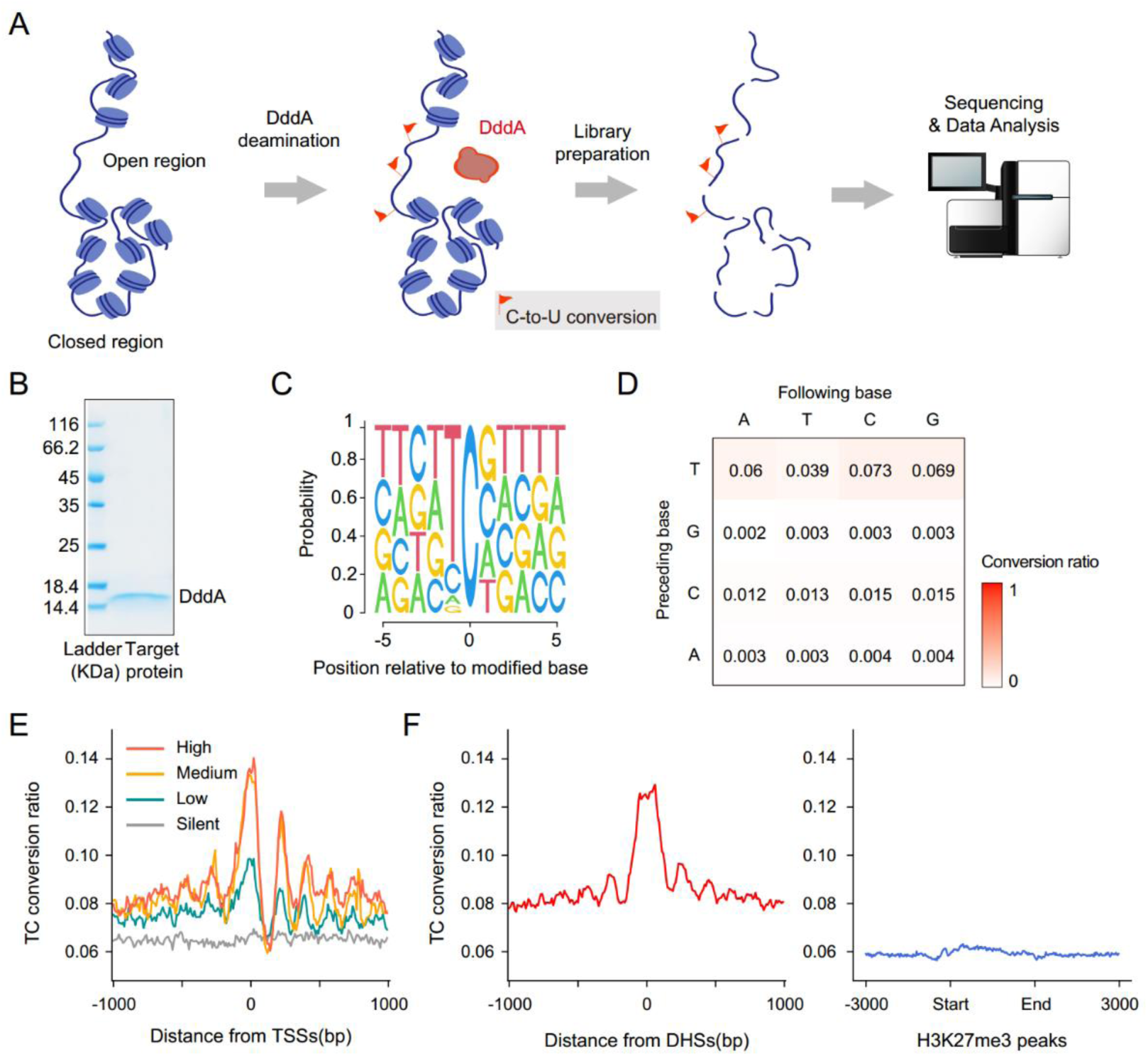
Workflow of ChrODIE for detecting open chromatin and the corresponding cytosine conversion of genomic DNA. (A) Schematic representation of the ChrODIE workflow. The double-stranded DNA deaminase DddA catalyzes cytosine (C) to uracil (U) conversion. After deamination, genomic DNA remains intact and is processed for library preparation, sequencing, and analysis. (B) Gel image of purified DddA. (C) Sequence logo of nucleotide frequency at positions relative to C nucleotides converted by DddA on genomic DNA. (D) DddA’s C to U conversion efficiency of ChrODIE on genomic DNA. The x-axis is the base following, and the y-axis base proceeding the cytidine. (E) Conversion ratios around transcription start sites (TSS) stratified by gene expression levels. Higher TC conversion ratios are observed near TSS, particularly in active genes. (F) Conversion ratios around DNase hypersensitive sites (DHS, left) and H3K27me3 repressed genes (right). The TC conversion ratio is high near DHS but low in H3K27me3-enriched regions.

C to U conversion by ChrODIE produces a genome-wide background of approximately 6% cytosine conversion (Fig. 1D), with conversion occurring preferably in open chromatin regions (Fig. 1E, F). Conversion ratios are strongly enriched near transcription start sites (TSS) of genes, with higher ratios for genes with elevated expression levels. Similarly, the conversion ratios are higher at DNase hypersensitive sites (DHS), but not at the TSS of repressed genes (Fig. 1F). These results indicate that ChrODIE deamination is enriched in regions of open chromatin.

The low level of cytosine conversion required by ChrODIE relative to methods like FOODIE allow a broad range of conversion ratios that can be tuned to maximize detection sensitivity of open chromatin regions. In the experiments described here, reads were typically 150bp (with next generation sequencing), with a positive read in an open chromatin region defined to have at least 20% conversion.

### Comparison with Other Open Chromatin Detection Methods

We compared ChrODIE to established methods for detection of open chromatin, including ATAC-seq and DNase-seq. Like ATAC-seq and DNase-seq, ChrODIE signal was clearly enriched around transcription start sites (TSSs) (Fig. 2A). ChrODIE produced a clear pattern of peaks (Fig. 2B) that were strongly concordant with ATAC-seq and DNase-seq and identified accessible regions with similar resolution.

**Figure 2.**
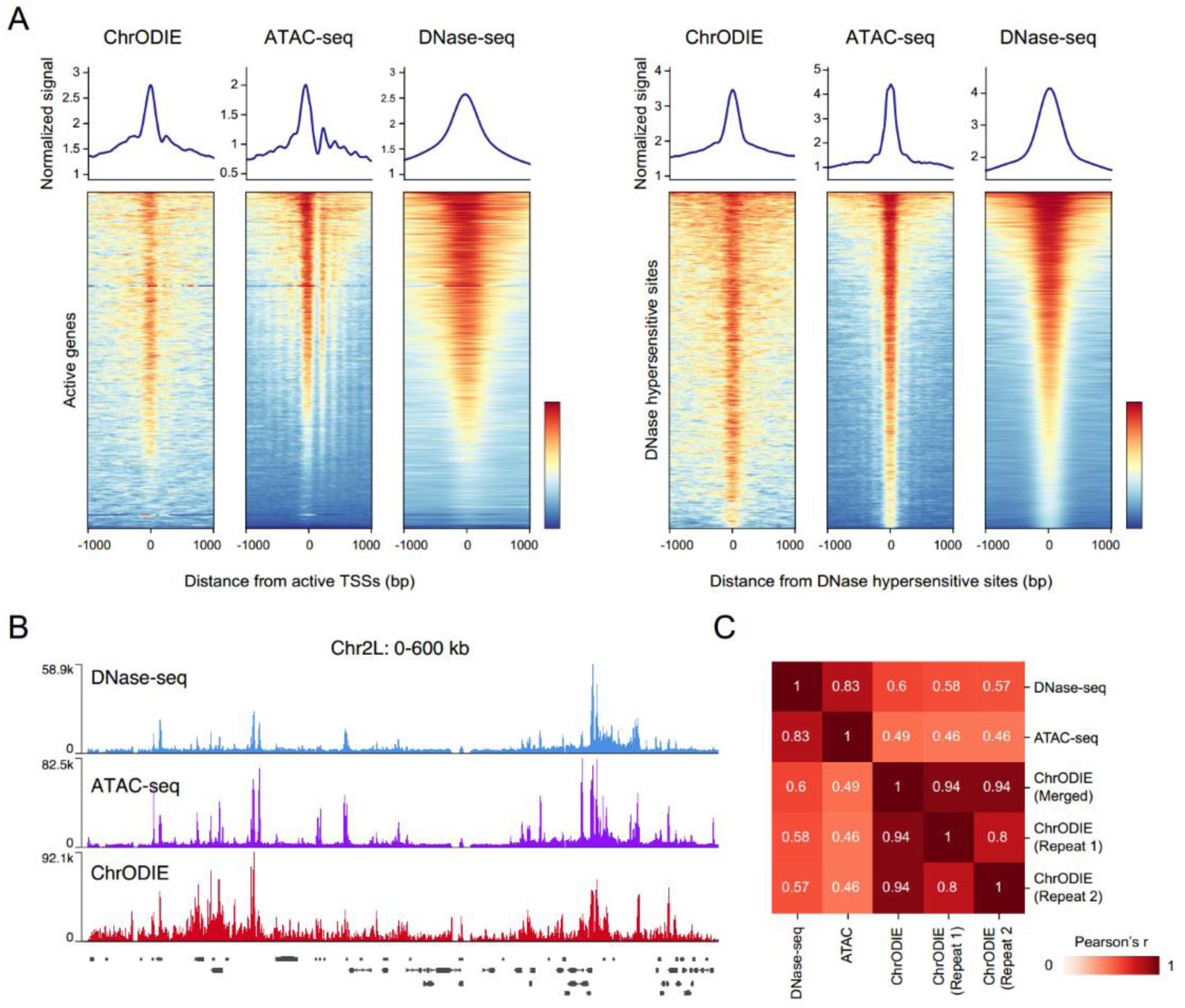
Comparison of ChrODIE with ATAC-seq and DNase-seq for chromatin accessibility detection. (A) Comparison of signals detected by ChrODIE, ATAC-seq, and DNase-seq around transcription start sites (TSS, left) and DNase hypersensitive sites (DHS, right). ChrODIE signals are enriched at both TSS and DHS regions, displaying patterns highly similar to those of ATAC-seq and DNase-seq. (B) Genome browser tracks for DNase-seq (blue), ATAC-seq (purple), and ChrODIE (red) signals across a representative genomic region (Chr2L: 0–600 kb). ChrODIE exhibits strong concordance with ATAC-seq and DNase-seq. (C) Heatmap of Pearson correlation coefficients between ChrODIE, ATAC-seq, and DNase-seq signals.

To further validate the performance of ChrODIE, we assessed its reproducibility and concordance with other chromatin accessibility methods through Pearson correlation analysis (Fig. 2C). ChrODIE demonstrated strong reproducibility across biological replicates (R=0.8), highlighting its robustness and reliability. ChrODIE exhibited moderate correlation with DNase-seq (R = 0.6) and ATAC-seq (R = 0.49), while DNase-seq and ATAC-seq exhibit high correlation with each other (R = 0.83). This suggests that ChrODIE captures open chromatin regions in common with ATAC-seq and DNase-seq, while potentially harboring method-specific patterns. A similar phenomenon has been reported in NOMe-seq, which has low correlation (R = 0.2) with ATAC-seq or DNase-seq (21).

### ANDIE detects histone modifications through antibody conjugation

When fused with protein A and protein G molecules (pAG) (Fig. 3A, 3B), dsDNA deaminase can be utilized for detecting histone modifications. ANDIE is based on antibody-directed deamination and begins with the incubation of permeabilized nuclei with antibodies specific to histone modifications (Fig. 3A). The binding of pAG-DddA to chromatin-bound histone modification antibodies enables site-specific deamination of cytosines proximal to the targeted histone modifications.

**Figure 3.**
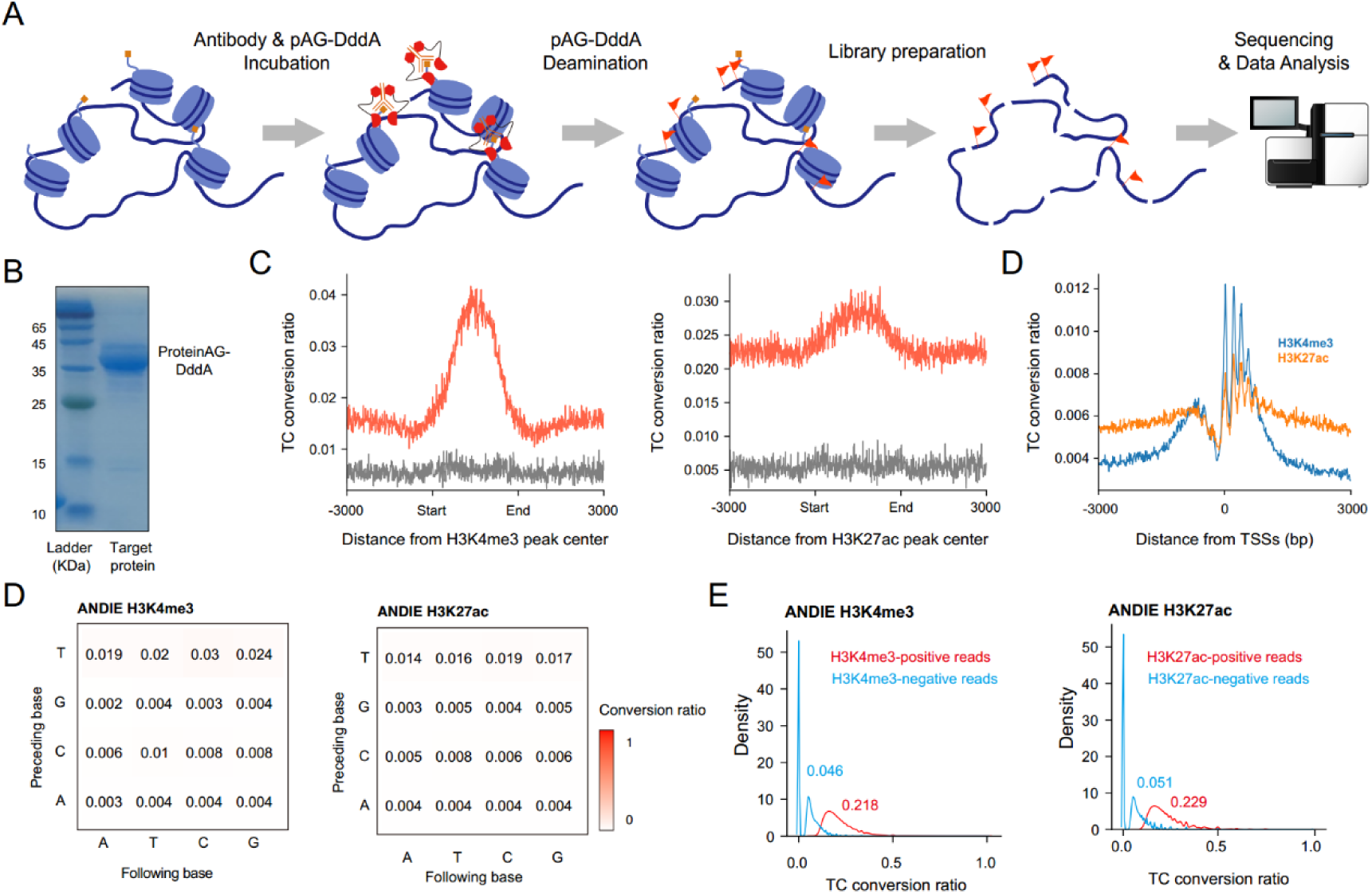
ANDIE workflow for profiling of histone modifications. (A) Lysed nuclei are incubated with antibodies conjugated to pAG-DddA, enabling targeted deamination of cytosines near antibody-bound regions. Library preparation and high-throughput sequencing determine conversion sites, allowing precise mapping of histone modifications. (B) Gel image of purified pAG-DddA. (C Conversion ratios (y-axis) around the peak centers of H3K4me3 (left panel) and H3K27ac (right panel). The x-axis shows distance from ChIP-seq peak centers. (D) Conversion ratios (y-axis) for H3K4me3 and H3K27ac modifications in TSS regions. The x-axis shows distance from TSS centers. (E) Genome-wide conversion ratios for H3K4me3 and H3K27ac modifications. Heatmaps represent deamination efficiencies across different DNA sequence contexts, with rows and columns corresponding to the preceding and following bases, respectively. (F) Density plots show the TC conversion ratios for ANDIE targeting H3K4me3 (left) and H3K27ac (right). Red curves represent positive reads, while blue curves represent negative reads. H3K4me3-positive reads exhibit a mean TC conversion ratio of 0.218 compared to 0.046 in negative reads (*P* < 0.0001, two-sided Mann-Whitney U rank test), and H3K27ac-positive reads show a mean ratio of 0.229 compared to 0.051 in negative reads (*P* < 0.0001, two-sided Mann-Whitney U rank test).

We applied ANDIE to acetylation of histone H3 on lysine 27 (H3K27ac), a marker of active enhancers, and trimethylation of histone H3 on lysine 4 (H3K4me3), which is commonly found at the promoters of actively transcribed genes. ANDIE with either H3K4me3 or H3K27ac antibodies showed conversion enrichment at ChIP-seq peak centers, with H3K4me3 showing sharp signals and H3K27ac displaying broader signals.

ANDIE with IgG control resulted in negligible background conversion (Fig. 3C). Signal enrichments were also observed at TSSs and show clear ∼150 bp signal periodicity, corresponding to the known location of positioned nucleosomes at TSS (Fig. 3D). These data demonstrate the specificity of ANDIE deamination.

Global TC conversion ratios of ANDIE for detection histone modifications (∼2%, Fig. 3E) are more than two-fold lower than that of ChrODIE (∼4%, Fig. 1D). Correspondingly, a positive read in a particular histone modification region was defined to have at least three converted TC sites within the paired-end 150bp NGS sequencing reads, with a minimum conversion ratio of 7% for H3K4me3 and 6% for H3K27ac (Fig. 3F). The mean TC conversion ratio of reads in histone modification-positive regions (∼20%) is about five-fold higher than comparable non-target regions (∼4%, with zero-conversion reads excluded).

### Comparison with Other Histone Modification Detection Methods

We compared the performance of ANDIE to established methods such as CUT&Tag and ChIP-seq. For the detection of H3K4me3, a mark enriched in promoter regions, all three methods showed consistent aggregate enrichment patterns when centered on active TSSs (Fig. 4A). We calculated the correlations between ANDIE, CUT&Tag, and ChIP-seq. H3K4me3 ANDIE replicates yielded correlation coefficients above 0.95, and values of above 0.8 with each of the other two methods (Fig. 4B). ANDIE for H3K27ac also exhibited high inter-replicate reproducibility and strong concordance with both CUT&Tag and ChIP-seq (Fig. 4B).

**Figure 4.**
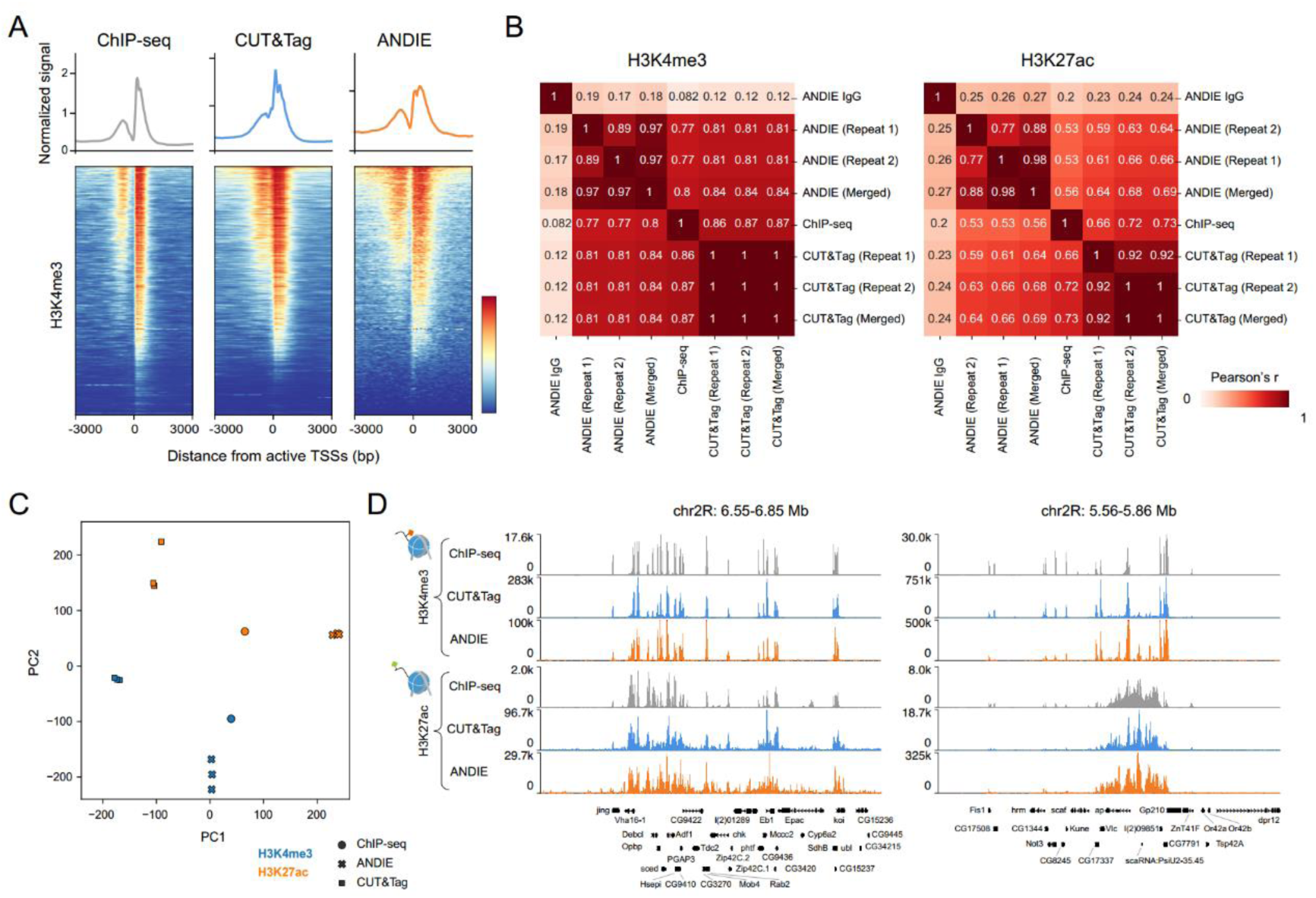
Comparison of ANDIE, CUT&Tag, and ChIP-seq for histone modification profiling. (A) Aggregate enrichment plots and heatmaps showing the distribution of H3K4me3 signals across transcription start sites (TSSs) for ANDIE, CUT&Tag, and ChIP-seq. All three methods display strong enrichment at TSSs. (B) Pearson correlation heatmaps for H3K4me3 (left) and H3K27ac (right), showing high inter-replicate correlations for ANDIE and strong concordance with CUT&Tag and ChIP-seq, with negligible correlations for the IgG control. (C) PCA plot illustrating the clustering of samples based on their genome-wide histone modification profiles, generated using ChIP-seq, CUT&Tag, and ANDIE. Each point represents a biological replicate for either H3K4me3 (blue) or H3K27ac (orange). Clear separation is observed between H3K4me3 and H3K27ac marks mapped by three different methods along the principal components. (D) Genome browser tracks of representative genomic loci for H3K4me3 (top) and H3K27ac (bottom). ANDIE accurately recapitulates the peaks observed with CUT&Tag and ChIP-seq.

While H3K4me3 and H3K27ac are generally associated with different regions of the genome, they are known to co-exist in highly active regulatory elements (22). We performed principal component analysis (PCA) to compare H3K4me3 and H3K27ac profiles generated by ANDIE, CUT&Tag, and ChIP-seq. Replicates for each method clustered tightly together, with a clear separation between H3K4me3 and H3K27ac marks along the principal components (Fig. 4C). We chose two genomic regions for visual comparison of histone mark localizations with ANDIE (Fig. 4D). On Chr2R (6.55-6.8 Mb), CUT&Tag and ChIP-seq produce nearly identical H3K4me3 and H3K27ac peaks, a pattern recapitulated by ANDIE with a strong signal-to-noise ratio. In contrast, nearby on Chr2R (5.56-5.86 Mb) CUT&Tag and ChIP-seq produced sharp H3K4me3 peaks but broad H3K27ac distributions, a pattern also recapitulated by ANDIE. These results demonstrate that ANDIE is an effective method for histone mark detection in diverse genome regions, with sensitivity and resolution comparable to existing CUT&Tag and ChIP-seq methods.

## Discussion

Chromatin features including nucleosome organization and histone modifications are critical for transcriptional regulation and regulation of other DNA-templated cellular processes. Mapping these chromatin features provides insight to underlying mechanisms of genome regulation. Epigenetic profiling with dsDNA deaminase achieves high reproducibility and results consistent with established techniques, supporting its utility as a reliable and complementary approach for genome-wide open chromatin or histone modification profiling. Further, deamination preserves genomic DNA integrity and enables detection of converted cytosines after amplification, which offers unique advantages for integration with long-read sequencing technologies like PacBio or Nanopore sequencing, even with few (or single) input cells, which cannot be achieved by current long-read techniques (Fiber-seq or DiMeLo-seq) that require a large number of input cells. This makes it particularly suitable for profiling chromatin dynamics in challenging or low-abundance samples, such as biopsies or progenitor cells. ANDIE and ChrODIE provide an opportunity to study long-range co-accessibility and chromatin interactions across multiple genomic regions in rare cell types. Combined with Hi-C, this method could provide insights into the interplay between 3D genome organization and chromatin accessibility. The integration of these technologies will enable more comprehensive analyses of gene regulation, chromatin structure, and chromatin interactions.

## Acknowledgments

This project is financially supported by Changping Laboratory, the Ministry of Science and Technology of China, Beijing Advanced Innovation Center for Genomics at Peking University and Peking-Tsinghua Center for Life Sciences.

## Materials and Methods

### Plasmid construction

The DddA expression plasmid was obtained from Genescript. To generate a pETDuet-1-based expression construct for DddA, the DddA encoding sequence: ATGGGCAGCAGCCATCACCATCATCACCACAGCCAGGATCCGGGTAGCTATGCGCTGGGTCCGTATCAGATCTCTGCTCCGCAGCTGCCGGCATATAACGGTCAGACTGTTGGTACTTTCTATTATGTTAACGATGCTGGCGGTTTAGAAAGCAAAGTTTTCAGCTCTGGTGGTCCGACCCCGTATCCGAACTATGCTAACGCTGGTCACGTTGAAGGTCAGTCTGCTCTGTTCATGCGTGATAACGGTATCTCTGAAGGTCTGGTTTTCCATAACAACCCGGAAGGTACCTGTGGTTTTTGTGTTAACATGACCGAAACCCTGCTGCCGGAAAACGCTAAAATGACCGTTGTTCCGCCGGAAGGTGCGATTCCGGTTAAACGTGGTGCTACCGGTGAAACCAA AGTTTTCACCGGTAACTCTAACTCTCCGAAATCTCCGACCAAAGGTGGTTGCTAA (of which the translated protein sequence is: MGSSHHHHHHSQDPGSYALGPYQISAPQLPAYNGQTVGTFYYVNDAGGLESKVFSSGGPTPYPNYANAGHVEGQSALFMRDNGISEGLVFHNNPEGTCGFCVNMTETLLPENAKMTVVPPEGAIPVKRGATGETKVFTGNSNSPKSPTKGGC.), is synthesized and cloned into the MCS-1 (NcoI and HindIII sites, retaining an N-terminal hexahistidine tag), and the immunity protein DddAI encoding sequence: ATGTATGCGGATGACTTTGACGGGGAAATTGAGATTGATGAAGTTGATAGCCTAGTTGAGTTTCTGAGCCGTCGTCCGGCGTTCGATGCGAACAACTTCGTTCTGACCTTCGAAGAAAGCGGCTTCCCGCAGCTGAACATCTTCGCGAAAAACGATATCGCGGTTGTTTACTACATGGATATCGGCGAAAACTTCGTTAGCAAAGGCAACAGCGCGAGCGGCGGCACCGAAAAATTCTACGAAAACAAACTGGGCGGCGAAGTTGATCTGAGCAAAGATTGCGTTGTTAGCAAAGAACAG ATGATCGAAGCGGCGAAACAGTTCTTCGCGACCAAACAGCGTCCGGAACAGCTGACCTGGAGCGAACTGTAA (of which the translated protein sequence is: MYADDFDGEIEIDEVDSLVEFLSRRPAFDANNFVLTFEESGFPQLNIFAKNDIAVVYYMDIGENFVSKGNSASGGTEKFYENKLGGEVDLSKDCVVSKEQMIEAAKQFFATKQRPEQLTWSEL.), is synthesized and cloned into the MCS-2 (NdeI and XhoI sites, removing an C-terminal S-tag). The Protein A-Protein G fused DddA expression plasmid was obtained from Genescript. To generate a pETDuet-1-based expression construct for pAG-DddA, the pAG-DddA encoding sequence: ATGGGCAGCAGCCATCACCATCATCACCACAGCCAGGATCCAACCATGATTACGCCAAGCTTAAAAGATGACCCAAGCCAAAGTGCTAACCTATTGTCAGAAGCTAAAAAGTTAAATGAATCTCAAGCACCGAAAGCGGATAACAAATTCAACAAAGAACAACAAAATGCTTTCTATGAAATCTTACATTTACCTAACTTAAACGAAGAACAACGCAATGGTTTCATCCAAAGCCTAAAAGATGACCCAAGCCAAAGCGCTAACCTTTTAGCAGAAGCTAAAAAGCTAAATGATGCTCAAGCACCAAAAGCTGACAACAAATTCAACAAAGAACAACAAAATGCTTTCTATGAAATTTTACATTTACCTAACTTAACTGAAGAACAACGTAACGGCTTCATCCAAAGCCTTAAAGACGATCCTTCAGTGAGCAAAGAAATTTTAGCAGAAGCTAAAAAGCTAAACGATGCTCAAGCACCAAAAACAACTTATAAATTAGTCATCAACGGGAAAACGCTGAAGGGTGAAACCACGACAGAGGCCGTAGATGCGGAGACAGCGGAGCGCCACTTTAAGCAATACGCGAATGATAACGGTGTAGACGGCGAGTGGACCTACGACGACGCGACAAAGACCTTTACCGTCACGGAGAAACCTGAGGTTATCGACGCGTCTGAGTTGACGCCAGCCGTAGGCTCTGGTGGTTCTAGCGGTGGTTCTTCTGGTTCCGGGGATCCGGGTAGCTATGCGCTGGGTCCGTATCAGATCTCTGCTCCGCAGCTGCCGGCATATAACGGTCAGACTGTTGGTACTTTCTATTATGTTAACGATGCTGGCGGTTTAGAAAGCAAAGTTTTCAGCTCTGGTGGTCCGACCCCGTATCCGAACTATGCTAACGCTGGTCACGTTGAAGGTCAGTCTGCTCTGTTCATGCGTGATAACGGTATCTCTGAAGGTCTGGTTTTCCATAACAACCCGGAAGGTACCTGTGGTTTTTGTGTTAACATGACCGAAACCCTGCTGCCGGAAAACGCTAAAATGACCGTTGTTCCGCCGGAAGGTGCGATTCCGGTTAAACGTGGTGCTACCGGTGAAACCAAAGT TTTCACCGGTAACTCTAACTCTCCGAAATCTCCGACCAAAGGTGGTTGCTAA (of which the translated protein sequence is: MGSSHHHHHHSQDPTMITPSLKDDPSQSANLLSEAKKLNESQAPKADNKFNKEQQNAFYEILHLPNLNEEQRNGFIQSLKDDPSQSANLLAEAKKLNDAQAPKADNKFNKEQQNAFYEILHLPNLTEEQRNGFIQSLKDDPSVSKEILAEAKKLNDAQAPKTTYKLVINGKTLKGETTTEAVDAETAERHFKQYANDNGVDGEWTYDDATKTFTVTEKPEVIDASELTPAVGSGGSSGGSSGSGDPGSYALGPYQISAPQLPAYNGQTVGTFYYVNDAGGLESKVFSSGGPTPYPNYANAGHVEGQSALFMRD NGISEGLVFHNNPEGTCGFCVNMTETLLPENAKMTVVPPEGAIPVKRGATGETKVFTGNSNSPKSPTKGGC.), is synthesized and cloned into the MCS-1 (NcoI and NotI sites, retaining an N-terminal hexahistidine tag), and the immunity protein DddAI encoding sequence: ATGTATGCGGATGACTTTGACGGGGAAATTGAGATTGATGAAGTTGATAGCCTAGTTGAGTTTCTGAGCCGTCGTCCGGCGTTCGATGCGAACAACTTCGTTCTGACCTTCGAAGAAAGCGGCTTCCCGCAGCTGAACATCTTCGCGAAAAACGATATCGCGGTTGTTTACTACATGGATATCGGCGAAAACTTCGTTAGCAAAGGCAACAGCGCGAGCGGCGGCACCGAAAAATTCTACGAAAACAAACTGGGCGGCGAAGTTGATCTGAGCAAAGATTGCGTTGTTAGCAAAGAACAG ATGATCGAAGCGGCGAAACAGTTCTTCGCGACCAAACAGCGTCCGGAACAGCTGACCTGGAGCGAACTGTAA (of which the translated protein sequence is: MYADDFDGEIEIDEVDSLVEFLSRRPAFDANNFVLTFEESGFPQLNIFAKNDIA VVYYMDIGENFVSKGNSASGGTEKFYENKLGGEVDLSKDCVVSKEQMIEAAKQFFATKQRPEQLTWSEL.), is synthesized and cloned into the MCS-2 (NdeI and XhoI sites, removing an C-terminal S-tag). The plasmids were transformed into *E. coli* strains DH5α BL21 and stored at −20 degrees.

### Bacterial strains and culture conditions

All *Escherichia coli* (*E. coli*) strains used in this study were grown in Lysogeny Broth (LB) at 37 °C or on LB medium solidified with agar (LBA, 1.5% w/v). The media was supplemented with ampicillin (100 μg per ml) or IPTG (0.5 mM) if it is required. *E. coli* strains DH5α and BL21 were used for plasmid maintenance and protein expression, respectively.

### Production of dsDNA deaminases

To purify the his-tagged deaminases in complex with their immunity protein, *E. coli* BL21 was used to inoculate 2 L of LB broth in a 1:100 dilution and cultured overnight. After the culture reached an OD600 of approximately 0.6, 0.5 mM IPTG was added, and the culture was incubated with shaking for 16 hours at 18°C. Bacterial cell pellets were collected by centrifugation at 4000 g for 30 minutes and then resuspended in 50 ml of lysis buffer (50 mM Tris-HCl pH 8.0, 500 mM NaCl, 10 mM imidazole, 1 mg/ml lysozyme, and a protease inhibitor cocktail). The cells were lysed by sonication (five pulses, 10 seconds each), and the supernatant was separated from the debris by centrifugation at 25,000 g for 30 minutes. The his-tagged deaminase-immunity protein complex was purified from the supernatant using a Nickel column and eluted with elution buffer (50 mM Tris-HCl pH 7.5, 300 mM NaCl, 30 mM imidazole, and 1 mM DTT). The eluted deaminase-immunity protein complex was denatured by adding 50 ml of 8 M urea denaturing buffer (50 mM Tris-HCl pH 7.5, 300 mM NaCl, and 1 mM DTT) and incubated for 16 hours at 4°C. The denatured proteins in the 8 M urea denaturing buffer were reloaded onto the Nickel column, which was then washed with 50 ml of the same buffer to remove any remaining immunity protein. Sequential washes were performed with denaturing buffer containing decreasing concentrations of urea (6 M, 4 M, 2 M, 1 M), followed by a final wash with buffer without urea. Deaminase bound to the column was eluted with 5 ml of elution buffer. The eluted deaminase was subjected to size-exclusion chromatography using fast protein liquid chromatography (FPLC) on a Superdex200 column (GE Healthcare) with sizing buffer (20 mM Tris-HCl pH 7.5, 200 mM NaCl, 1 mM DTT, and 5% (w/v) glycerol). The purity of the eluted deaminase was assessed by SDS-PAGE gel stained with Coomassie blue (Fig. S1B), and the protein was stored at −80°C.

### Cell culture and preparation

Drosophila Schneider 2 (S2) cells were gifted from Dr. Yan Song at Peking University. S2 cells were cultured in Schneider’s Drosophila Medium (Gibco, cat. no. 21720024) supplemented with 10% fetal bovine serum (FBS) (Cellmax, cat. no. SA301.02) and 1% Penicillin-Streptomycin (Pen-Strep) (Gibco, Thermo Fisher Scientific, cat. no. 15140122) at 25°C without CO2. Upon use, S2 cells were collected by centrifuge at 300g for 5 min and resuspended in 1×PBS. When fixation (0.5% PFA) is needed, 3 million S2 cells were resuspended in 500µL 0.5% PFA for 10 minutes. Fixed cells were collected by centrifuge at 600g for 5 min, resuspend in 300µL 1% BSA/PBS for 5 min to quench PFA, and lastly washed three times with 1% BSA-PBS.

### Open chromatin labelling through deamination (ChrODIE)

A total of 500,000 0.5% PFA fixed S2 cells were lysed with 150µL cold lysis buffer (10 mM Tris-HCl, pH 7.5, 50 mM NaCl, 0.5 mM DTT and 0.1% IGEPAL-CA630) by incubating on ice for 5 min, then collected by centrifuge at 600g for 5 min. Lysed nuclei were resuspended in 50µL 1× deamination buffer (10 mM Tris-HCl, pH 7.5, 50 mM NaCl, 0.5 mM DTT) containing a titration of DddA enzyme. The incubation reaction was performed at 37°C for 30 minutes with gentle rotating. After deamination, nuclei were washed twice with PBS. After that, genomic DNA extraction.

A negative control of nuclei without deamination can be set before genomic DNA extraction. For genomic DNA extraction, the nuclei were collected by centrifuge at 600g for 5 min, and resuspend in 20µL 0.1%SDS-PBS with 2µL proteinase K (P8107S, NEB). After overnight incubation at 55°C with lid-heat on, the cleanup reaction was performed with a Zymo DNA Clean and Concentrator-5 Kit (D4014, Zymo). Library preparation with eluted genomic DNA can be performed using TruePrep DNA Library Prep Kit (TD501, Vazyme) following the manual. In general, a genome-wide TC conversion ratio at around 6% after deamination by DddA at certain titration is ideal for open chromatin labelling.

### Histone modification detected by CUT&Tag

CUT&Tag was performed with 0.5% PFA fixed S2 cells using Hyperactive Universal CUT&Tag Assay Kit for Illumina Pro (Vazyme, TD904-01) following the manual. H3K4me3 antibody (Abcam, ab213224) and H3K27ac antibody (Abcam, ab4729) were used in CUT&Tag experiments. For each group, 3×10^5 S2 cells were used.

### Histone modification detected by ANDIE

Prepare Wash buffer by mixing 50 µL 1 M HEPES pH7.5, 75 µL 5 M NaCl, 31.5 µL 0.1 M spermidine, and add 1X Roche Complete Protease Inhibitor (50 µL from one tablet in 1 mL), 12.5 µL 20% BSA, and 2.281 mL dH2O. Prepare Antibody buffer by mixing 1 µL 0.5 M EDTA, 2.5 µL 10% Tween-20 (final 0.1%) and 246.5 µL Wash buffer. Prepare High-salt buffer by mixing 1 µL 0.5 M EDTA, 15 µL 5 M NaCl and 500 µL Wash buffer. Prepare Incubation buffer by mixing 2 µL 1M Tris pH 7.5, 2 µL 5M NaCl, 1 µL 0.1M DTT, 95 µL dH2O.

S2 cells were rinsed with PBS and resuspend in PBS for cell counting. To mildly fix the cells, resuspend approximately 3×10^6 S2 cells in 500 µL of 0.5% PFA, rotate for 5 minutes, or gently invert to ensure thorough contact between the fixative and the cells. Subsequently, add 300 µL of 1% BSA/PBS, mix well, and centrifuge at 500×g for 4 minutes at 4°C, discarding the supernatant. Resuspend the cells in 300 µL of 1% BSA/PBS, incubate with rotation for 5 minutes, then resuspend in 200 µL of PBS and count the cells. Transfer 5×10^5 S2 cells to a PCR tube, centrifuge at 800×g for 5 minutes at 4 ° C, and discard the supernatant. Add 1 µL of 10% NP-40 (final concentration 0.1%) to 100 µL of Wash buffer. Resuspend the cells with 100 µL of Wash buffer containing 0.1% NP-40 in the PCR tube, and incubate on ice for 5 minutes. Immediately centrifuge at 800×g for 5 minutes at 4°C and discard the supernatant. Wash the cells once with 100 µL of Wash buffer, centrifuge at 800×g for 5 minutes at 4°C, and remove the supernatant. Next, add 1 µL of antibody (with one IgG control group (CST, CST3678S), one H3K4me3 experimental group (Abcam, ab213224), and one H3K27ac experimental group (Abcam, ab4729)) to 50 µL of ice-cold Antibody buffer, mix thoroughly, and resuspend the pellet. Incubate with rotation at room temperature for 2 hours. After incubation, centrifuge the S2 cells at 300×g for 5 minutes, wash once gently with 100 µL of chilled Antibody buffer. Resuspend the pellet, centrifuge the S2 cells at 300×g for 5 minutes at 4°C and discard the supernatant. Gently resuspend the pellet in 50 µL of ice-cold High-salt buffer containing 2.5 µL of pAG-DddA, and incubate on a rotator for 1 hour. Incubate at low temperature (4°C) with rotating, to enable the binding of pAG-DddA to antibody but prevent deamination. Centrifuge at 300×g for 5 minutes and discard the supernatant. Wash the pellet twice with 150 µL of chilled high-salt buffer, gently pipetting about 10 times. After resuspending, centrifuge the S2 cells at 300×g for 5 minutes at 4°C and discard the supernatant. Gently dissolve the pellet in 100 µL of Incubation buffer and incubate at 37°C for 1 hour, which allows for antibody directed deamination. Finally, centrifuge the S2 cells at 800×g for 5 minutes, discard the supernatant, and add 100 µL of 0.2% SDS-PBS (prepared by adding 2 µL of 10% SDS to 98 µL of PBS) and 10 µL of proteinase K (P8107S, NEB). Incubate overnight at 55°C. After the incubation, extract and purify the genomic DNA using the Zymo Research D4010 Genomic DNA Clean & Concentrator®-10 and measure the DNA concentration using a Qubit fluorometer.

Library preparation with eluted genomic DNA can be performed using TruePrep DNA Library Prep Kit (TD501, Vazyme) following the manual. In general, a genome-wide TC conversion ratio at around 2% after antibody directed deamination targeting specific histone modification is ideal for detection.

### CUT&Tag data analysis

Raw sequencing reads were aligned to the dm6 reference genome using bowtie2 (version 2.3.4.3) with “ --end-to-end --very-sensitive --no-mixed --no-discordant --phred33 -I 10 -X 700”. PCR duplicates were removed using the Picard tool. The bigwig files were generated using bamCoverage (deeptools, version 3.5.1).

### ChrODIE and ANDIE analysis

Raw sequencing reads were aligned to the dm6 reference genome using biscuit (version 1.5.0). PCR duplicates were then counted and removed using dupsifter (version 1.0.0). The TC conversion sites were extracted using biscuit tools with default parameters.

For ChrODIE analysis, to obtain open chromatin signals, we discarded reads with low mapping quality scores (MAPQ<20) and used a custom Python script to selected reads with >20% TC-converted sites. For ANDIE analysis, to obtain signals near specific histone modifications, we discarded reads with low mapping quality scores (MAPQ<20) and selected reads with at least 3 TC-converted sites. The selection of the cutoff value is based on testing different cutoff values. Current two criteria in use have shown the highest correlation with the reference data.

The bigwig files were generated from above filtered BAM files using bamCoverage for downstream analysis and visualization. The average TC conversion ratios and coverage around TSSs, DHS and peaks were calculated using computeMatrix.

### Published data

*Drosophila* S2 published bulk RNA-seq, ATAC-seq and ChIP-seq data of Drosophila S2 cells were downloaded from the Gene Expression Omnibus (ATAC-seq: GSE103177; RNA-seq: GSE145320; H3K4me3: GSM1017409 and GSM2175508; H3K27ac: GSM1017404 and GSM2521732).

Published DNase-seq data were taken from the online flybrain resource website (http://compbio.med.harvard.edu/flychromatin/data.html). CrossMap and liftOver were used to convert genomic coordinates from dm3 to dm6.

